# Hypoxia regulate developmental coronary angiogenesis potentially through VEGFR2- and SOX17-mediated signaling

**DOI:** 10.1101/2023.08.16.553531

**Authors:** Halie E. Vitali, Bryce Kuschel, Chhiring Sherpa, Brendan W. Jones, Nisha Jacob, Syeda A. Madiha, Sam Elliott, Eddie Dziennik, Lily Kreun, Cora Conatser, Bhupal P. Bhetwal, Bikram Sharma

**Affiliations:** Department of Biology, Ball State University, Muncie, IN 47306; Division of Biomedical Sciences, College of Osteopathic Medicine, Marian University, Indianapolis, IN 46222

**Keywords:** Hif-1alpha, Endothelial cells, Sinus venosus, Endocardium, APJ

## Abstract

**Background:** Coronary vessels in embryonic mouse heart arises from multiple progenitor population including sinus venosus (SV), endocardium, and proepicardium. ELA/APJ signaling is shown to regulate coronary growth from SV pathway within the subepicardium, whereas VEGF-A/VEGF-R2 pathways is implicated to regulate coronary growth from endocardium pathway. Our previous study show hypoxia as a potential signaling cue to stimulate overall coronary growth and expansion within the myocardium. However, the role of hypoxia and its downstream signaling pathways in the regulation of coronary vessel development is not known. In this study, we investigated the role of hypoxia in coronary vessel development and have identified SOX17- and VEGF-R2-mediated signaling as a potential downstream pathway of hypoxia in the regulation of coronary vessel development.

**Results:** We show that hypoxia gain-of-function in the myocardium through upregulation of HIF-1α disrupts the normal pattern of coronary angiogenesis in developing mouse hearts and displays phenotype that is reminiscent of accelerated coronary growth. We show that VEGF-R2 expression is increased in coronary endothelial cells under hypoxia gain-of-function *in vivo* and *in vitro*. Furthermore, we show that SOX17 expression is upregulated in developing mouse heart under hypoxia gain-of-function conditions, whereas SOX17 expression is repressed under hypoxia loss-of-function conditions. Furthermore, our results show that SOX17 loss-of-function disrupts normal pattern of coronary growth.

**Conclusion:** Collectively, our data provide strong phenotypic evidence to show that hypoxia might regulate coronary growth in the developing mouse heart potentially through VEGF-R2- and SOX17-mediated downstream signaling pathways.

## Introduction

Defects in coronary artery are involved in the pathogenesis of fetal disease such as myocardial infarction, which results in heart failure. Coronary artery related diseases are one of the leading causes of death worldwide [1]. Despite its clinical significance, there is no cure to remedy damaged coronary arteries. By harnessing the mechanistic details of early progenitor pathways of coronary vessel development in the embryo, we can then reactivate these pathways to repair and regenerate damaged coronary arteries in adults. Our previous study implicated myocardial hypoxia as the potential stimulant of coronary vessel growth in the developing mouse heart [2]. However, the role of hypoxia and its downstream angiogenic pathways in the regulation of coronary vessel development has not been investigated. In this study, we investigated whether hypoxia regulates the developmental coronary angiogenesis and identified its potential downstream signaling pathways.

During development, coronary vessels are formed through a stepwise angiogenic program: first immature vascular plexus is formed by endothelial cells (ECs) and then it remodels into mature coronary arteries and veins [3-5]. ECs also line the luminal wall of coronary arteries and veins, which are covered by smooth muscle layers and outer adventitial layers. An important aspect of coronary vessel development in mice is the origin of coronary ECs from multiple progenitor pathways [4-6]. Studies in mice show that the source of ECs that form the initial capillary network arise from multiple progenitor sources namely proepicardium [7], sinus venosus [8], and endocardium [9]. The contributions from multiple sources combine to establish the entirety of coronary vascular tree. Progenitors from SV and endocardium are the two major contributors of coronary ECs [5]. Coronary vessels from SV first grow within the subepicardium and later enters myocardium, whereas coronary vessels from endocardium grow directly into the myocardium. Our previous study suggests that these two progenitor pathways are under the control of different molecular programs. For example, the sinus venosus (SV) progenitor pathway requires ELA/APJ [2] and VEGF-C [5] to grow in the subepicardium, whereas the endocardial pathway appears to respond to signals such as VEGF-A in the myocardium. Our previous study implicated hypoxia as a potential signaling cue to stimulate coronary growth within the myocardium from its dual progenitor pathways [2, 9]. However, the role of hypoxia and its detail molecular mechanisms in the regulation of coronary vessel development is not known.

A constant supply of oxygen is needed to support organogenesis. Tissue hypoxia develops when oxygen levels drop too low due to the lack of vessels or poor vascular permeability [10]. There are certain genes that are upregulated when cells are in the hypoxic environment. One of the genes that is upregulated is known as hypoxic-inducible factor-1α (*HIF-1*α), which encodes for the transcription factor HIF-1α. HIF-1α is upregulated when oxygen levels are too low in tissues and it activates angiogenic genes such as *Vegf-A* and *Cxcl12* [11]. By induction of angiogenic genes, hypoxia can stimulate the growth of blood vessels in hypoxic tissues. Under normal oxygen levels, HIF-1α undergoes prolyl hydroxylation; during this, a protein called von Hippel-Lindau (VHL) will bind and induce ubiquitin-mediated degradation of HIF-1α [12]. This indicates that VHL is a negative regulator of HIF-1α protein levels. Under hypoxic conditions, there is a decrease in the ability of prolyl hydroxylase to hydroxylate the HIF-1α subunit. This will result in stabilization and heterodimerization of HIF-1α [13]. HIF-1α was found to be abundant in the regions of developing mouse heart where coronary vessels arise from its progenitor [2]. A study reported a developmental defect in the mouse heart due to VEGF-A signaling deficiency [9, 14], indicating that the poor heart development in VEGF-A mutant was due to insufficient coronary vessel growth in the myocardium. Therefore, it is possible that hypoxia/HIF-1α/VEGF-A signaling might stimulate coronary angiogenesis in the myocardium. However, there is no direct evidence suggesting that myocardial hypoxia stimulates coronary angiogenesis from its progenitor sources. Determining the effect of both gain-of-function and loss-of-function of hypoxia is needed to confirm the role of hypoxia in the myocardial growth of coronary vessels.

Sex Determining Region Y-Box 17 (SOX17) is a transcription factor that is shown to determine the fate of progenitor cells in retinal arteries of mice [15]. A lack of SOX17 in arterial ECs is lethal *in utero*, suggesting that SOX17 is important for arterial differentiation [15]. In addition to its function in arterial differentiation, several studies show that SOX17 promotes sprouting angiogenesis. In addition to matured arteries, SOX17 is also found to be expressed by the coronary capillary ECs [2]. In tumors, SOX17 is expressed in tumor ECs and is shown to promote tumor angiogenesis by upregulating VEGFR2 expression in sprouting ECs [16]. SOX17 deletion in tumor ECs normalized tumor vessels and inhibited tumor growth and progression. SOX17 is also shown to be important for endothelial regeneration upon inflammation induced vascular injury [17]. In this injury model, SOX17 increased endothelial proliferation via upregulation of Cyclin E1. Endotoxemia upregulates hypoxia inducible factor alpha, which in turn transcriptionally activates SOX17. SOX17-OE *in vivo* is shown to enhance lung endothelial regeneration. Furthermore, SOX17 is known to promote sprouting angiogenesis partly by inducing a tip cell differentiation of ECs by activating endothelial tip cell markers such VEGFR-2, APELIN, ESM1, ANG2, PDGFb, and DLL4 [18]. Interestingly, we found that SOX17 is expressed by activated coronary progenitor cells that give rise to coronary vessels. Despite its expression in coronary vessels and its precursor cells, the exact role of SOX17 in coronary angiogenesis is unknown. Here, we investigated whether SOX17 is important for coronary angiogenesis and functions through hypoxia induced signaling axis in the regulation of coronary vessel formation.

## Methods and Materials

### Mice Used

All experimental procedures involving animals were conducted in accordance with the Institutional Animal Care and Use Committee (IACUC) of Ball State University. The following mice were purchased from the Jackson Laboratory and used in this study: Mef2cCre (stock 025556), VHL ^flox/flox^ (stock 012933), and CD-1 (stock 002962).

### Mouse Breeding for Timed Pregnancy

Male and female mice were bred to produce timed pregnancy. Mice were maintained under standard laboratory conditions (temperature: 25 ± 2º C, humidity: 60 ± 5%, 12-hour dark/light cycle) and fed with a standard laboratory diet and water. Breeding was maintained until the embryos reach the appropriate age for analysis. To determine embryonic age, the morning of vaginal plug detection was designated as embryonic day E0.5 and vaginal plugs were checked two times a day, once in the morning and once in the evening.

### Embryo and heart isolation from timed pregnancies

Female mice with timed pregnancies were sacrificed via CO_2_ euthanization. Following CO_2_ euthanization, cervical dislocation was performed to ensure death. Upon cervical dislocation, pregnant mice were subjected to embryo harvest. For embryo harvest, a female was placed on the dissection pad on her dorsal side, and her belly was sprayed with 70% ethanol to sterilize and wet the skin surface. Then, a T-shaped incision was made with scissors to open the pelvic cavity and expose the uterine horn carrying the embryos. The uterine horn was then dissected and placed in ice-cold, sterile 1X phosphate-buffered saline (PBS) (Fisher Scientific, catalog# SH3025602). Using forceps, embryos were removed from the uterine horn by peeling off tissues. For histological analysis, embryos were fixated using 4% PFA in 1X PBS. To ensure perfusion of fixative, embryos were decapitated, and incisions were made around the belly. For other analyses such as in vitro explant assays and RNA isolation and analysis, embryos were not fixed but immediately subjected to heart removal and its processing for experiments. Hearts were removed from the thoracic cavity using forceps, as described in Large et al. (2020). Isolated hearts were then subjected to further processing and analysis as needed.

### Ventricular explant culture

Ventricular explant culture was performed similarly as previously described in Large et. al 2020 with some modification. Briefly, isolated hearts were placed in a petri dish full of sterile 1X PBS. Lobes were cut out from each lung using fine forceps. Hearts were oriented on their dorsal side. Both atriums, sinus venosus, and adjacent tissue that surrounds the sinus venosus were removed using fine forceps. The outflow (aorta and pulmonary trunk) was removed from heart and remaining ventricles containing the endocardium were transferred to a new petri dish containing ice-cold sterile 1X PBS. Isolated sinus venosus and whole ventricles were kept on ice.

Explants were cultured in a PET culture inserts (MCEP24H48, Millicell) placed in a 24-well plate. Explants were cultured in endothelial growth media-2 complete (EGM-2). Cultures were placed in 37°C, 21% oxygen incubator for normoxia and in a 37°C, 5% oxygen incubator for hypoxia. Cultures were routinely observed in an inverted light microscope to ensure explants exhibit contractile beating and are attached to the bottom of the membrane. On day 2, complete media was removed and replaced by endothelial basal media (EBM) + 2% FBS. 24-well plates were placed back into their specific incubator and were grown for 5 days. After 5-day culture, media was removed, and explants were washed with 1X PBS at room temperature. Explants were fixed with 4% PFA in 4°C for 20 minutes while rocking. 4% PFA was removed from cultures and the cultures were washed with 1X PBS at least 3 times for 10 minutes before the samples were subjected to further analysis.

### Immunostaining and Imaging

Fixed explant heart cultures were subjected to immunostaining. Immunostaining was performed either in PCR tubes (for whole heart staining), 24-well plates (for the explant cultures), or on slides (for cryosection samples). Primary and secondary antibodies were diluted in blocking solution (5% donkey serum, 0.5% PBT). The primary antibodies that were used are: DACH1 (1:500, Cat:10914-AP, Protein Tech), a nuclear marker for coronary endothelial cells, Erg 1/2/3 (1:1000, Cat: ab92513, abcam), an endothelial cell nucleus antibody, VEGFR2 (1:100 Cat: AF644, R&D Systems), VE-Cadherin (1:125, Cat:550548, BD Pharmingen), an endothelial cell membrane marker, HIF-1α (1:100, Cat: NB100-479, Novus Biologicals), SOX17 (1:100, R&D systems). Samples were rocked overnight at 4°C. The next day, the samples were washed in 1X PBS (for cryosection samples) and in .5% PBT (for whole mount and explant cultures) for 30 mins (3 times 10 min each wash). Following the wash steps, samples were subjected to secondary antibody staining. The secondary antibodies that were used are: Donkey Anti-Rat Alexa Fluor 488 (1:250, Cat: A21208, Invitrogen), Donkey Anti-Rabbit Alexa Fluor 555 (1:250, Cat: A31572, Invitrogen), and Donkey Anti-Goat Alexa Flour 488 (1:250, Cat: A21081, Invitrogen). Samples were rocked overnight in secondary antibody at 4°C. After secondary antibody incubation, samples were washed with appropriate wash buffers (like primary antibody) for 30 minutes. Whole mount samples were incubated in mounting media. Explant cultures were removed from the inserts and mounted in slides with Vectashield containing DAPI (Cat: H-1200, Vector Labs) and sealed in coverslip for explant cultures as previously described. Slides and hearts were imaged using Zeiss LSM 5 Pascal Inverted microscope.

### qRT-PCR analysis

Total RNA from the whole heart samples were isolated using the Qiagen RNeasy Mini Kit (Fisher Scientific, catalog# 74104) strictly following the manufacturer’s instructions. The total concentration of isolated RNA was measured by using a NanoDrop One device (Fisher Scientific, catalog #13-400-519). The RNA was then subjected gDNA removal using DNase I enzyme mix following the manufacturer’s instruction. cDNA is obtained by performing reverse transcriptase reaction using BIO-RAD iScript reagent as per their recommendation. SYBR green based real time qPCR reaction was prepared using So-Advanced Universal SYBR green Supermix reagents (Bio-Rad, catalog #172-5271) in a 96-well plate and analyzed using the Bio-Rad CFX Opus RT-qPCR equipment. The reactions were run in triplicate for each condition. The expression of each target genes were normalized with GAPDH as a reference gene. The expression data was analyzed using Bio-Rad CFX Opus software.

### Statistical analyses

A non-parametric unpaired *T-test* with Welch’s correction was performed to compare differences between the two samples groups using Prism 9 (GraphPad). Areas of coronary outgrowth was calculated using ImageJ/Fiji by subtracting area occupied by the original explant from total area occupied by the explant after culture. Density of Erg 1/2/3+ cells is calculated by counting Erg 1/2/3+ cells in four field of view (FOV) of equal area per explant. The statistical significance was determined as p < 0.05.

## Results

### Upregulation of HIF-1α in the myocardium disrupted normal pattern of coronary angiogenesis in developing mouse hearts

To determine the role of hypoxia in coronary vessel development *in vivo*, we sought to use a mouse model in which HIF-1α is upregulated in the myocardium by deleting von hippel lindau (*VHL)*. To knockout *VHL*, we utilized Cre-LoxP based gene editing system using Mef2cCre and *VHL flox/flox* mouse lines. In the *Cre+; VHL* ^*fl/fl*^ hearts, we wanted to confirm whether HIF-1α was indeed upregulated in the myocardium. To show this, we stained e13.5 embryonic hearts, isolated from control and *Cre+; VHL* ^*fl/fl* (^VHL cKO) embryos, with HIF-1α antibody and co-stained with VE-cadherin to visualize coronary vessels in the myocardium. We performed whole mount confocal imaging and analyzed the images. HIF-1α levels were high in the VHL cKO (Cre+; VHL ^fl/fl^) hearts compared to the control (Cre-; VHL ^fl/fl^) in the myocardium of the heart (**Figure 1, HIF-1**α **panel**). Our analysis of coronary plexus in the cKO hearts show that the coronary ECs are unable to form normal tubular capillary plexus but instead appear to be in a highly disorganized clump of cells compared to normal capillary plexus in the control (**Figure 1, VE-cadherin panel**). In addition, we also observed blood island formation (white arrowhead) in the cKOs compared to control hearts, which is one of the characteristics of active endocardial angiogenesis observed under hypoxia.

**Figure 1.**
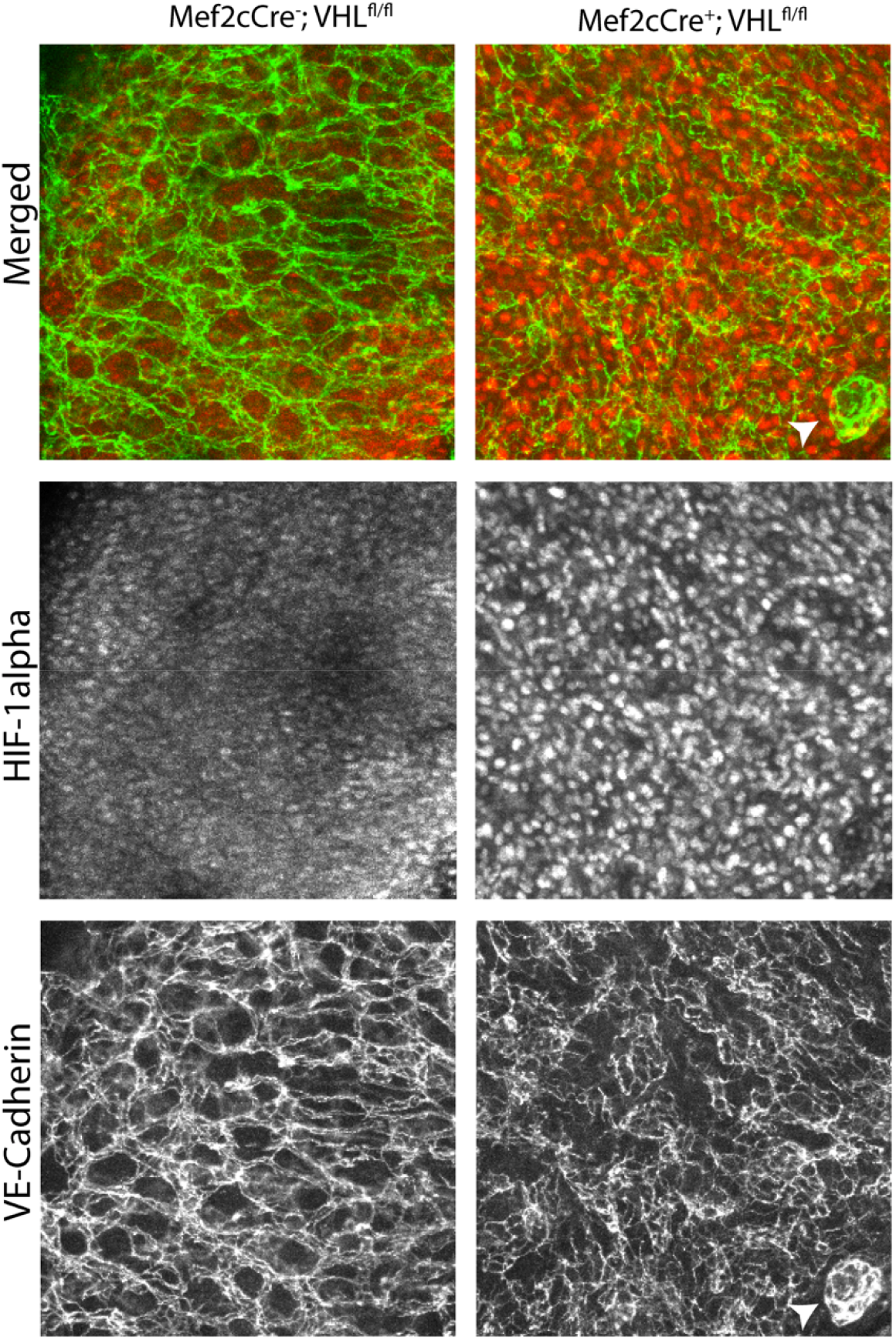
Upregulation of Hif-1alpha in VHL cKO hearts disrupts the normal pattern of coronary vessel growth. (A) Confocal imaging to label coronary endothelial cells (VE-Cadherin, green) and tissue hypoxia (HIF-1α, red) in the myocardium of control (left column) and myocardium specific VHL KO (right column) of E13.5 mouse hearts. High level of HIF-1α expression (HIF-1α row) is observed in VHL cKO hearts compared to control hearts. Abnormal coronary growth pattern is observed in cKO hearts compared to control hearts. Contol, N= 3; VHL cKO, N = 3.

### Upregulation of HIF-1α in the myocardium accelerated coronary growth in the myocardium

To determine the role of hypoxia in coronary vessel growth *in vivo*, we analyzed coronary plexus in e13.5 hearts and compared between the control and cKO groups. Previous studies have shown compartmental contribution of coronary vessel growth from sinus venosus and endocardium progenitors [5]. It has been shown that sinus venosus-derived coronary plexus initially grew onto the dorsal surface and invaded into the myocardium from outside-in growth. On the other hand, endocardium progenitors began its growth from inside-out into the septal myocardium and out into the myocardium of free walls. Therefore, sinus venosus-derived vessels dominantly feed the dorsal side of the heart and endocardium-derived vessels feed the mid ventral region of the heart. We analyzed coronary growth patterns in these spatial locations to understand the effect of myocardial hypoxia in coronary growth from the sinus venosus and endocardium.

On the mid-ventral side, where coronary growth is dominantly from the endocardium, we observed that coronary plexuses on the surface were well-formed into web-like structures in the cKO hearts where hypoxia was activated. In contrast, the well-formed capillary plexuses were less in the control hearts. Instead, we observed that endothelial blood islands were much more abundant in the control hearts compared to the cKO hearts (**Figure 2A, see arrowhead**). When we analyzed coronary vessel growth in the intramyocardial regions on the ventral side (endocardial pathway), we observed similar phenotypes where there were more well-formed coronary plexuses in the hypoxic hearts compared to control group (**Figure 2B**). Quantification of DACH1+ endothelial cells over many ventral surface hearts showed similar coronary endothelial density between hypoxic cKO and control hearts. Quantification of coronary vessel branching on the ventral surface show slightly increased coronary vessel branching in hypoxic cKO hearts, but it was not statistically significant. Similarly, coronary vessel density in the ventral intramyocardium was similar between the control hearts and hypoxic cKO hearts.

**Figure 2.**
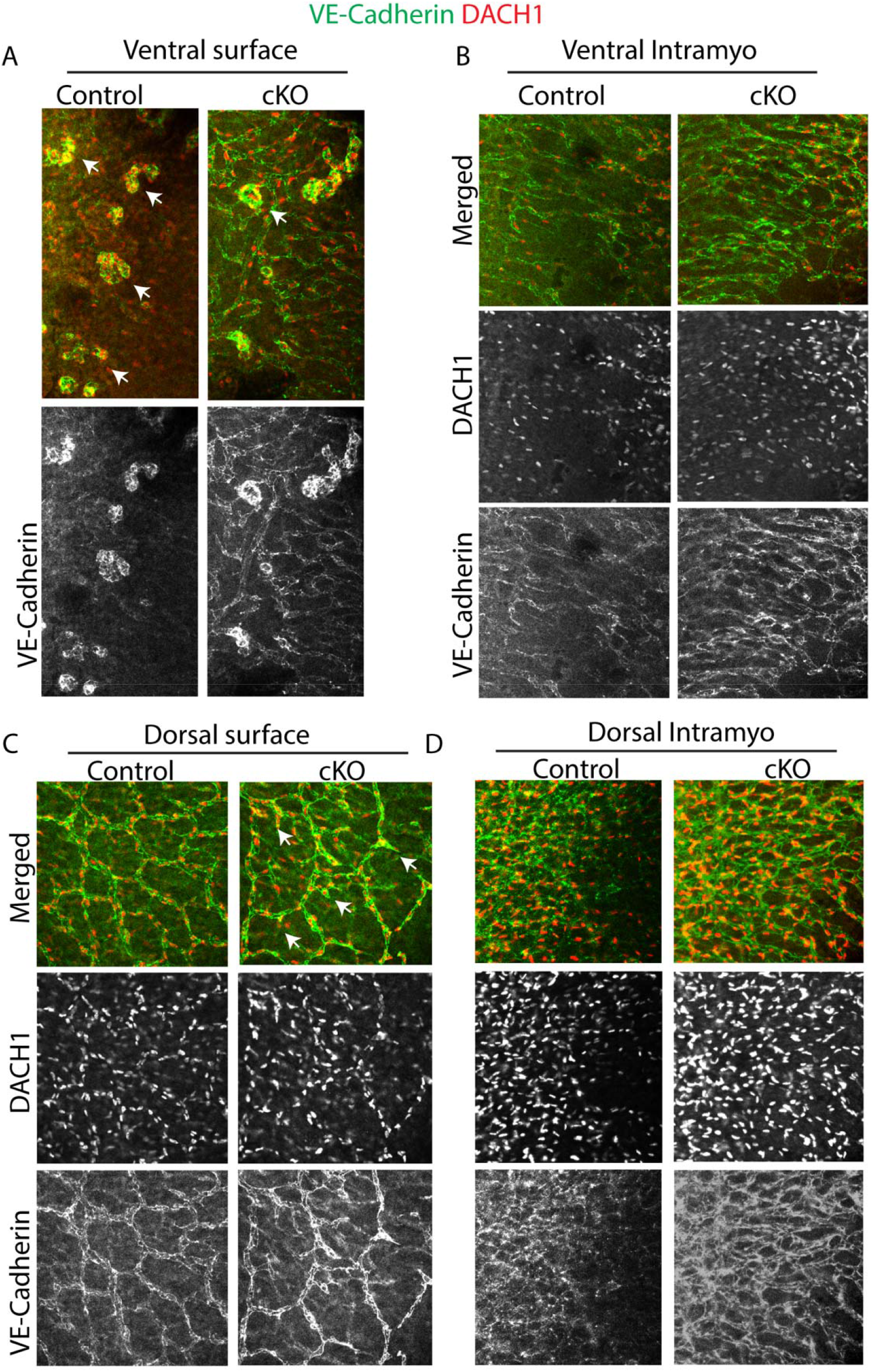
Normal pattern of coronary growth is disrupted in VHL cKO hypoxic hearts compared to control. Whole-mount immunostaining of E13.5 control and VHL cKO hearts immunostained for VE-cadherin (endothelial cells surface, green) and DACH1 (coronary endothelial cell nuclei, red) showing ventral surface (A), ventral intramyocardium (B), dorsal surface (C), and dorsal intramyocardium (D) regions of the heart. Arrowheads in A pointing as blood island-like structures and in C pointing at the disrupted capillary web-like structures or discontinued capillary network. Contol, N= 7; VHL cKO, N = 4.

To analyze the effect of myocardial hypoxia on the sinus venosus pathway of coronary angiogenesis, we analyzed coronary vessel growth on the dorsal surface where coronary vessels initially grow from sinus venosus and eventually invade into the myocardium. On the dorsal surface, we observed broken capillary plexuses in the hypoxic cKO hearts compared to control (Figure 2C, see arrowheads). On the control hearts, there were well formed capillary plexuses that spread over the dorsal surface. The capillary plexuses in the hypoxic cKO hearts were found to have significantly reduced vessel branch points (**Figure 2C**). However, coronary vessel density in control were slightly increased compared to hypoxic hearts (**Figure 2C**). Coronary plexuses were found to be well-formed and more advanced in the dorsal intramyocardium region of the hypoxic cKO hearts compared to control, although coronary endothelial cell density was comparable between the two groups (**Figure 2D**).

### Hypoxia regulate coronary angiogenesis potentially through VEGFR2 signaling

To determine the role of hypoxia on endocardium-derived coronary angiogenesis, we subjected ventricles of E11.5 embryonic hearts (age before the coronary growth in the myocardium begins) to explant cultures under hypoxic (5% oxygen) and normoxic (21% oxygen) conditions.

Ventricular explant cultures showed robust coronary growth under hypoxia compared to normoxia. 5-day culture displayed increase in coronary expansion since the area of coronary coverage was significantly higher in hypoxic conditions compared to normoxia (**Figure 3A**). Furthermore, we observed robust VEGFR2 expression in endothelial cells under hypoxic conditions compared to normoxia. In the normoxic culture, VEGFR2 expression was restricted to endothelial cells on the migrating front, whereas VEGFR2 expression was ubiquitously expressed by endothelial cells in all areas of explant (**Figure 3A, boxed region**). In addition, the sprouts in the migrating fronts of hypoxic explants displayed pointed cells with filopodia-like extensions, whereas the cells were blunted and less pointed in the normoxic explants (**Figure 3A, Red boxed region, white arrows**). Quantification of coronary expansion (area covered) by active sprouts showed significant increase in hypoxic conditions compared to control groups (**Figure 3A’**). Quantification of Erg 1/2/3+ endothelial cells over many explants showed slightly increase coronary endothelial density in hypoxic conditions but was not statistically significant (**Figure 3 A’’**).

**Figure 3.**
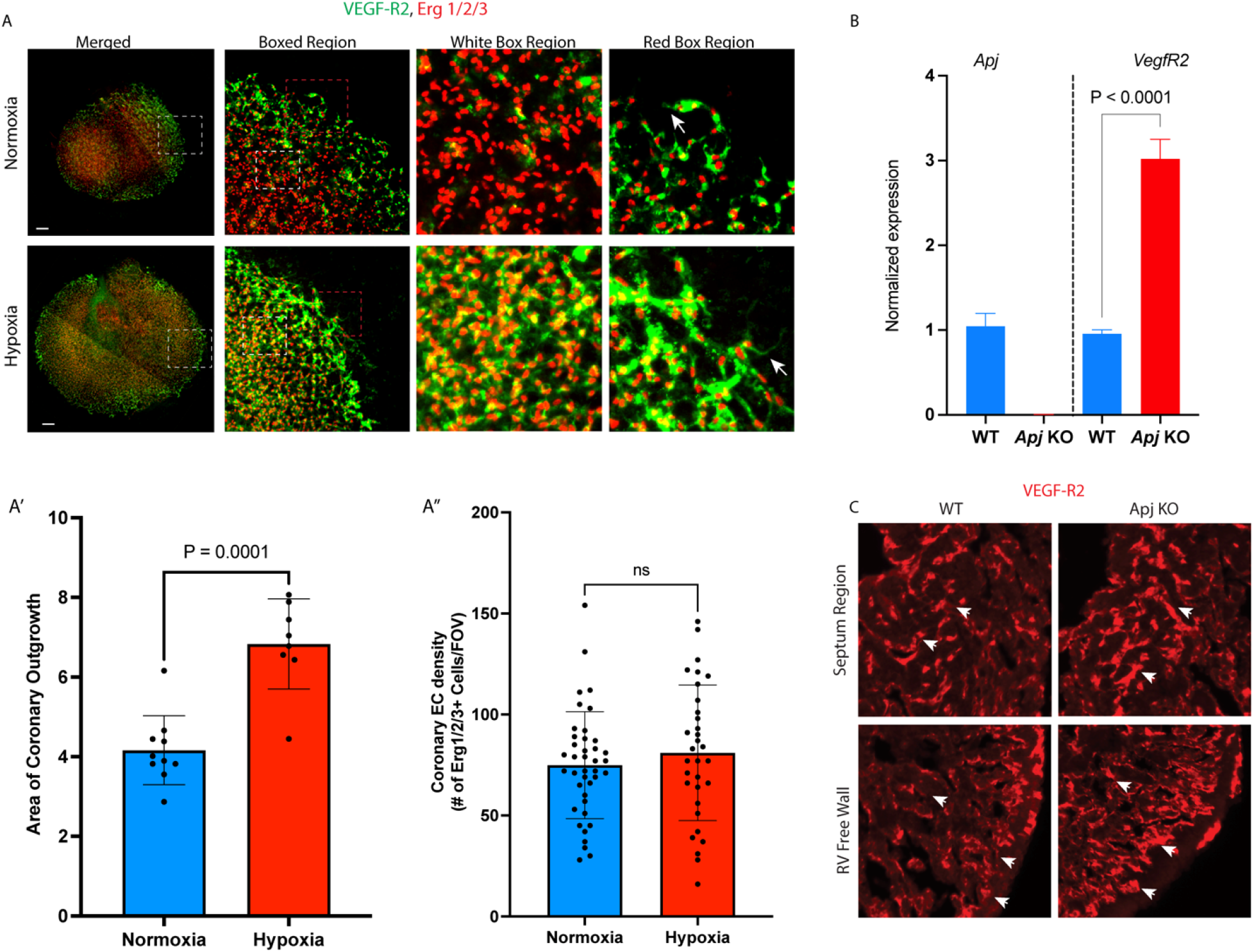
Hypoxia stimulate coronary growth expansion and increases VEGF-R2 expression in coronary endothelial cells. A) Whole-mount immunostaining of the E13.5 ventricular 5-day explant culture immunolabelled for VEGF-R2 (endothelial cell surface receptor for VEGF-A signaling) and Erg 1/2/3 (endothelial cell nuclei) show increased radial expansion of coronary growth in explants cultured under hypoxia compared to explants cultured in normoxia. Boxed regions show high magnification views of outlined regions. White arrows pointing at tip cell at the migrating front of coronary sprouts showing filopodial extensions. A’ and A”) Quantification of coronary outgrowth (A’) and coronary endothelial density (A”). B) qPCR analysis showing Apj (left panel) and Vegf-R2 (Right panel) RNA expression comparison in wildtype (WT) and Apj knockout (KO) hearts. C) Immunostaining of cryosection samples of E14.5 wildtype (WT) and Apj knockout (KO) mouse hearts immunolabeled for VEGF-R2 (in red). Arrowheads pointing at VEGF-R2 expression in WT and KO samples showing differences in VEGF-R2 expression levels. Scale bars, 50 μm (A). Bar graph represents Mean±SD. N = 10, Normoxia; N = 8, Hypoxia.

Next, to determine whether hypoxia stimulated coronary growth through a VEGFR2-mediated signaling in coronary endothelial cells, we analyzed VEGFR2 expression in control versus hypoxia gain-of-function hearts. In our previous study [2], we showed that the APJ KO hearts were highly hypoxic since Hif-1alpha expression was significantly increased in the KO hearts compared to control. We utilized APJ KO hearts as a hypoxia gain-of-function model to determine whether VEGFR2 expression in coronary endothelial cells is upregulated under hypoxia. To test this, we analyzed VEGF-R2 expression in APJ KO hearts using qPCR. As expected, qPCR analysis detected robust APJ expression in WT hearts but did not detect any APJ expression in APJ KO hearts (**Figure 3A**). Furthermore, qPCR analysis showed almost twofold increase in VEGF-R2 RNA expression levels in APJ KO hearts compared to WT hearts (**Figure 3B**). Similarly, immunostaining analysis of VEGF-R2 expression in the septum and free ventricular wall of embryonic heart sections revealed stronger VEGF-R2 staining in the hypoxic APJ KO hearts compared to relatively weaker VEGF-R2 staining in WT hearts (**Figure 3C, arrowheads**). Overall, our results show increased VEGF-R2 expression in angiogenic coronary endothelial cells under hypoxic conditions compared to normoxia.

### Hypoxia regulate coronary angiogenesis potentially through Sox17 mediated signaling

Our previous study indicated that Sox17 might be activated by hypoxia in coronary endothelial cells since we observed robust Sox17 expression in hypoxic region of developing mouse hearts [2]. To determine whether Sox17 is involved in the regulation of coronary angiogenesis through hypoxia mediated signaling, we first analyzed Sox17 expression in hypoxia gain-of-function and loss-of-function models of developing mouse hearts. We observed the upregulation of Sox17 expression in highly angiogenic septal regions of the developing mouse hearts in VHL cKO (hypoxia gain-of-function) hearts compared to control hearts (**Figure 4A**). Quantification of Sox17+ cells within the septal region revealed almost threefold increase in cKO hearts compared to control. Furthermore, we observed severe downregulation of Sox17 expression in hif-1alpha cKO (hypoxia loss-of-function) hearts compared to control hearts (**Figure 4B**). Our results strongly suggest that Sox17 is a target of hypoxia in coronary endothelial cells. To further establish whether Sox17 is involved in the regulation of coronary angiogenesis, we analyzed the pattern of coronary growth in control versus Sox17 cKO hearts, where Sox17 is depleted from endocardial progenitor population using *BmxCreER* deletor line coupled with Sox17 flox mouse lines. We observed that the normal coronary growth pattern is disrupted in the cKO hearts compared to control hearts. Well-formed capillary sprouts were consistently found at this stage in the control hearts, whereas immature, more rounded endothelial blood island-like patches were observed in the cKO hearts (**Figure 4C, see VE-Cadherin panel**). The rounded endothelial patches reflects a phenotype reminiscent of early stages of endocardial angiogenesis [2, 5, 8], suggesting that the Sox17 cKO resulted in a delayed coronary angiogenesis phenotype. Overall, our results show strong evidence that the coronary angiogenesis is regulated by hypoxia mediated Sox17 activation in coronary endothelial progenitors.

**Figure 4.**
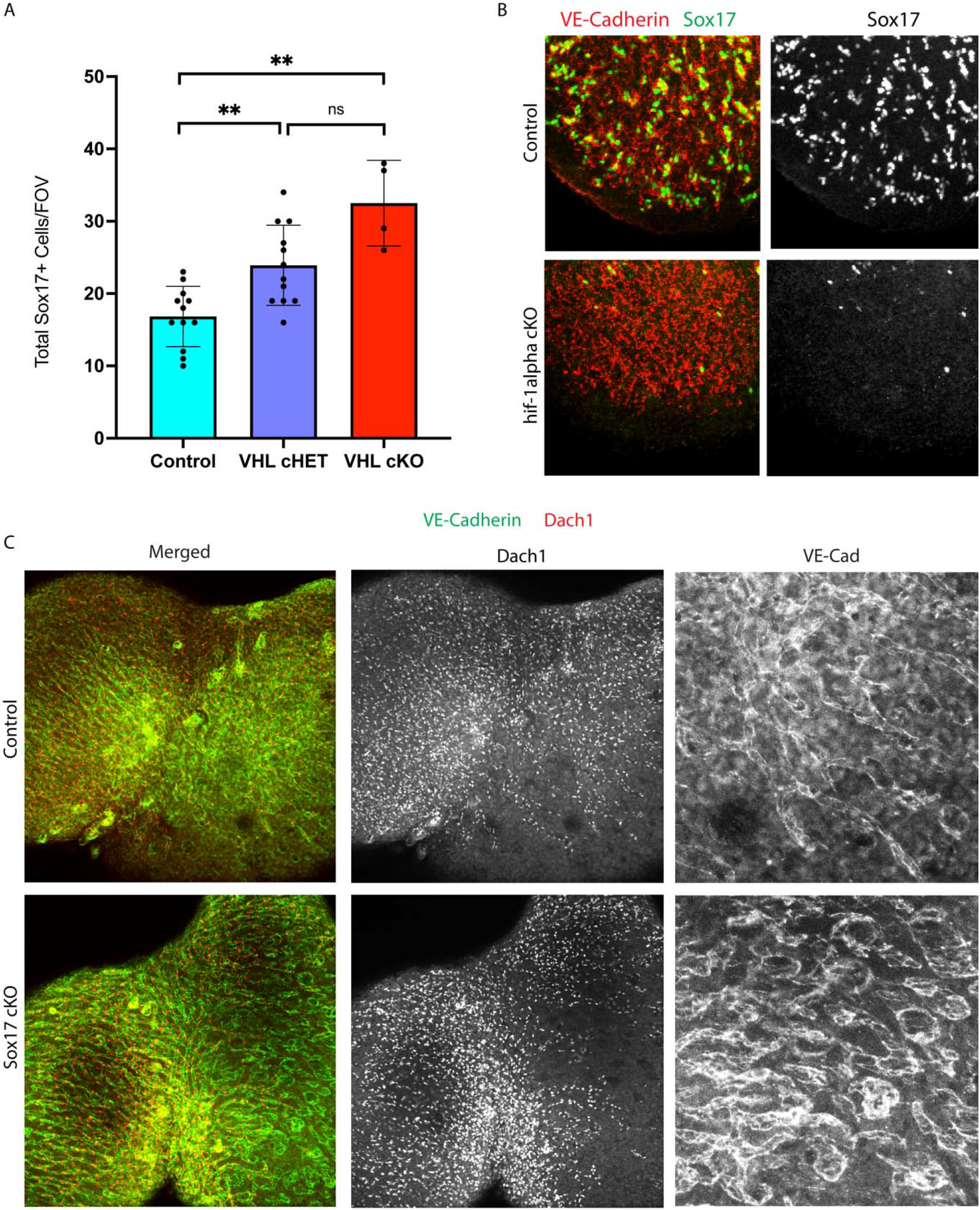
Hypoxia stimulate Sox17 expression and Sox17 regulate coronary growth. A) Bar graph show quantification of Sox17+ cells in various field of view within the septal region of the control, VHL cHET, and VHL cKO hearts analyzed by immunostaining of cryosection samples. B) Whole-mount confocal imaging of E12.5 control and hif-1alpha deficient hearts immunolabelled for VE-Cadherin (endothelial cells) and Sox17. C) Whole-mount immunostaining of the ventral aspect of E16.5 control and Sox17 cKO hearts immunolabelled for VE-Cadherin (endothelial cells) and Dach1 (coronary endothelial cells) show that normal pattern of coronary growth is disrupted in cKO with delayed coronary growth phenotype. A) Control = 3, VHL cHET = 3, VHL cKO = 1. B) Control = 4, Hif1alpha-cKO = 3. C) Control = 3, Sox17 cKO = 2. ** indicate *P* < .01; ns indicate not significant.

## Discussion

It is well known that the coronary vessels in the developing mouse heart grows from multiple progenitor pathways including the sinus venous and the endocardium [4]. While the sinus venosus pathway grow coronary vessels from epicardial surface into the myocardium, the endocardium pathways grow coronary vessels directly into the myocardium [5]. Epicardial cues such as VEGF-C and ELABELA is known to stimulate coronary vessels on the epicardial surface, but the signaling cues that stimulate coronary growth in the myocardium is not fully understood [2, 5]. VEGA/VEGF-R2 signaling is implicated as one of the signaling mechanisms in the myocardium but what activates VEGF-R2 signaling in the coronary endothelial cells/progenitor cells is not clear [19]. This study investigated hypoxia as a potential upstream signal that activates downstream angiogenic pathways including VEGF-A/VEGF-R2 and Sox17 mediated signaling to promote coronary growth in the myocardium. As the heart grows and adds layers of myocardium, the heart reaches a point where the luminal blood flow is insufficient to perfuse the thick myocardium. At this point, myocardium becomes hypoxic. We predicted that myocardial hypoxia stimulates coronary growth in the myocardium. Surprisingly, this possibility has not yet been tested directly. Here, we conducted experiments using hypoxia gain-of-function and loss-of-function mouse models *in vivo* and explant tissue cultures models *in vitro* to show that hypoxia stimulates coronary angiogenesis potentially through VEGF-R2- and Sox17-mediated downstream signaling pathways.

To determine how myocardial hypoxia impacts coronary vessel growth, we utilized *in vivo* mouse models where the hypoxia inducible factor, HIF-1α, is either upregulated or downregulated specifically in myocardium. HIF-1α is a transcription factor that is activated in tissues in response to hypoxia and is known to activate pro-angiogenic genes such as VEGFA and CxCL12, which then stimulate new blood vessel formation by angiogenesis to vascularize the hypoxic tissue. Hypoxic response will be activated by stabilizing HIF-1α in the myocardium by deleting VHL. VHL, in the presence of oxygen, degrade HIF-1α in tissues. In the absence of VHL, Hif1-alpha becomes stable and provides tissues a hypoxic response as if the tissues are in hypoxia. To gain hypoxic function, we deleted VHL specifically in the myocardium by crossing myocardium specific mouse Cre lines Mef2cCre with VHL floxed mouse lines. We show that HIF-1α is upregulated in the myocardium when VHL is floxed out using Mef2cCre+ line (**Figure 1, HIF-1alpha panel**). We observed abnormal capillary plexus phenotypes in hypoxic VHL cKO hearts that failed to form web-like capillary network as see in control groups. This phenotype is somewhat reminiscent of highly angiogenic coronary endothelial cells potentially under active proliferation stage that resulted in clumping of cells rather than fully differentiated endothelial cells forming capillary plexus (**Figure 1, VE-Cadherin Panel**). Collectively, our data suggested that upregulation of Hif-1alpha in the myocardium disrupted normal pattern of coronary growth potentially due to increased coronary endothelial cell stimulation through activation of hypoxia inducible downstream signaling cues such as VEGF-A/VEGF-R2 signaling.

We performed detail analysis of coronary phenotype both in the endocardium-derived coronary region in the ventral aspect of the heart as well as the sinus venous-derived coronary region on the dorsal aspect of the heart. Previous study reported that the endocardium derived vessel grow initially into the mid-ventral region of the myocardium extending from the septal region of the heart [5, 8]. Therefore, we analyzed coronary growth around the mid-ventral aspect of the heart on the surface as well as into the myocardium. Our phenotypic analysis on the ventral surface of the developing heart show that the hypoxia gain-of-function in VHL cKO hearts accelerated angiogenic activity from the endocardium. We observed more robust capillary network formation with few blood islands in the VHL cKO hearts compared to more primitive coronary growth phenotypes with fewer well-formed capillary networks but more abundant blood islands in the control hearts (**Figure 2A**). Blood islands are hallmarks of early-stage coronary growth from endocardium. Consistently, we also observed more developed coronary vessels in the myocardium of cKO hearts compared to control hearts in the mid-ventral region (**Figure 2B**). On the dorsal aspect, we observed well-formed and continuous coronary plexus on the surface of control hearts, but we observed disrupted and discontinuous capillary plexus with broken branches in the cKO hearts (**Figure 2C**). We predict that this phenotype is due to more dominant angiogenic signal in the myocardium due to hypoxia gain-of-function that accelerated inward growth (invasion) of surface coronary vessels. As expected, we found the increased vascular density in the myocardium of cKO hearts compared to control hearts (**Figure 2D**). Overall, our results show that hypoxia gain-of-function in the myocardium accelerated coronary growth into the myocardium from the sinus venosus- and endocardium-derived pathways.

Next, we investigated potential downstream signaling pathways mediated by hypoxia that is involved in the regulation of coronary growth. VEGF-R2 signaling has been previously implicated as a signaling mechanism that stimulate coronary growth in the myocardium [19]. We assayed whether hypoxia stimulate coronary angiogenesis in VEGF-R2 dependent manner. To investigate this, we performed an *in vitro* coronary angiogenesis assay using previously described ventricular explant culture assay. Results from the explant culture showed that hypoxia stimulated coronary growth as we observed increased radial outgrowth of coronary sprouts in hypoxic culture conditions compared to normoxia (**Figure 3A**). More importantly, we observed more robust VEGF-R2 expression in coronary endothelial cells of hypoxic explants compared to normoxia (**Figure 3A**). This result provides a strong connection that increased coronary angiogenesis under hypoxic condition is potentially due to increased VEGF-A/VEGF-R2 signaling. Furthermore, we also observed active angiogenic endothelial phenotype with apparent filopodia-like extensions in the sprouting fronts of hypoxia explants compared to more blunted endothelial phenotype in normoxia. Overall, our result provide a strong correlation between hypoxia, active coronary angiogenesis, and VEGF-R2 expression, which collectively suggest that hypoxia potentially stimulate coronary growth via VEGF-A/VEGF-R2 mediated signaling. Our results is consistent with reports from previous study showing VEGF-A/VEGF-R2 signaling accelerated coronary angiogenesis [19]. To further substantiate this possibility, we assayed VEGF-R2 expression in APJ knockout hearts, a known *in vivo* model with hypoxia gain-of-function in the myocardium. qPCR and immunostaining analysis showed increased VEGF-R2 expression in the APJ KO hearts compared to WT suggesting that VEGF-A/VEGF-R2 signaling in coronary endothelial cells stimulate coronary angiogenesis in the hypoxic myocardium (**Figure 3B**). This provide additional evidence to support hypoxia/VEGF-R2 mediated signaling axis for coronary vessel growth during embryonic mouse heart development.

In our previous study [2], we found the expression of SOX17, a transcription factor, in the endocardial coronary progenitors within the endocardium-myocardium border of the avascular and hypoxic region of the heart. We also observed that the SOX17 expressing endocardial progenitors sprouted in the avascular myocardial regions of the heart. This suggested to us that hypoxia may activate SOX17 in the endocardial progenitors to prime these cells to undergo coronary angiogenesis. To test this, we first needed to establish whether SOX17 is activated in the endocardium-derived coronary endothelial cells due to hypoxic cues in the myocardium. In this study, we performed experiments to directly test whether the expression of SOX17 is changed due to hypoxia gain-of-function or loss-of-function conditions *in vivo*. Immunostaining data revealed that Sox17 expression is indeed dependent upon hypoxic conditions. We found a positive correlation between hypoxia and SOX17 expression. In other words, when hypoxic condition is increased in VHL cKO hearts, we observed increased number of SOX17 expressing cells within the septal region of the heart, region of the developing heart known to have active coronary angiogenesis from endocardial progenitors, compared to control hearts (**Figure 4A**). Similarly, when hypoxic condition is depleted in Hif-1alpha cKO hearts, we observed dramatic decrease in SOX17 expressing cells within the myocardium of E12.5 hearts compared to its control group (**Figure 4B**). This result clearly show that SOX17 expression is dependent upon hypoxic condition of the myocardium. Next, we wanted to test whether SOX17 is important for coronary vessel growth. When we depleted SOX17 from endocardial progenitors using *BmxCreER* deletor line [20], we observed phenotype that is reminiscent of delayed coronary growth since the we observed immature coronary vessels with blood-island like structures in the cKO hearts compared to well refined coronary capillary plexus in the control hearts (**Figure 4C**). This provided a strong phenotypic data that the function of SOX17 is critical for proper formation/patterning of coronary vessel growth in the myocardium of the developing mouse heart.

Overall, we have provided strong phenotypic evidence to show that hypoxia is a critical signaling cue to drive coronary vessel growth in the myocardium of the developing mouse heart. We also show that hypoxia activates VEGF-R2 and SOX17, which are important in the regulation of coronary angiogenesis. Taken together, our study provides data to implicate hypoxia as an important signaling cue in the developing myocardium to stimulate coronary vessel growth potentially through activation of VEGF-A/VEGF-R2 signaling and SOX17 mediated signaling mechanisms. It is yet to determine whether VEGF-R2 and SOX17 work independently of one another or within the same signaling axis downstream of hypoxia in the regulation of coronary vessel growth in the developing mouse heart.

